# Deciphering active prophages from metagenomes

**DOI:** 10.1101/2021.01.29.428894

**Authors:** Kristopher Kieft, Karthik Anantharaman

## Abstract

Temperate phages (prophages) are ubiquitous in nature and persist as dormant components of host cells (lysogenic stage) before activating and lysing the host (lytic stage). Actively replicating prophages contribute to central community processes, such as enabling bacterial virulence, manipulating biogeochemical cycling, and driving microbial community diversification. Recent advances in sequencing technology have allowed for the identification and characterization of diverse phages, yet no approaches currently exist for identifying if a prophage has activated. Here, we present PropagAtE (Prophage Activity Estimator), an automated software tool for estimating if a prophage is in the lytic or lysogenic stage of infection. PropagAtE uses statistical analyses of prophage-to-host read coverage ratios to decipher actively replicating prophages, irrespective of whether prophages were induced or spontaneously activated. We demonstrate that PropagAtE is fast, accurate and sensitive, regardless of sequencing depth. Application of PropagAtE to prophages from 348 complex metagenomes from human gut, murine gut and soil environments identified distinct spatial and temporal prophage activation signatures, with the highest proportion of active prophages in murine gut samples. Among the soil habitats evaluated (bog, fen and palsa), we identified unique populations of Myxococcales, Acetobacteraceae and Acidimicrobiaceae prophages to be active in fen, palsa and bog habitats, respectively. Within the human gut, 11 prophage populations, some encoding the sulfur metabolism gene *cysH* or a *rhuM*-like virulence factor, were consistently present over time but not active. Overall, PropagAtE will facilitate accurate representations of viruses in microbiomes by associating prophages with their active roles in shaping microbial communities in nature.

## INTRODUCTION

Bacteriophages (phages) are pervasive entities that are ubiquitous on Earth. Phages drive evolutionary adaptation and diversification of microorganisms, play critical roles in global nutrient cycles and can directly impact human health (Breitbart and Rohwer 2005; Wommack and Colwell 2000; Gobler et al. 1997; Jiao et al. 2010; Fuhrman 1999; Wilhelm and Suttle 1999; Norman et al. 2015; Barr et al. 2013). Phages can be organized into two categories according to how they infect a host cell: lytic and temperate. Temperate phages are those that have the ability to integrate their dsDNA genome into their bacterial host and can be identified in nearly half of all cultivated bacteria (Touchon et al. 2016). These integrated prophage sequences can co-exist with the host cell in a lysogenic stage in which virions are not produced. During host genome replication the prophage sequence is likewise replicated in a one-to-one ratio. Given host-dependent or environmental cues such as DNA damage or nutrient stressors, or spontaneous activation, the prophage can enter a lytic stage to produce virions and lyse the host (Howard-Varona et al. 2017; Cochran et al. 1998; Casjens and Hendrix 2015; Carrolo et al. 2010; Binnenkade et al. 2014; Feiner et al. 2015). On the other hand, lytic phages are those that directly enter the lytic stage upon infection with no mechanism for integration and dormancy.

Prophages can affect their host and surrounding microbial communities in both the “dormant” lysogenic stage as well as in the “active” lytic stage. In the dormant stage, prophages can impose physiological changes on the host by altering gene expression patterns, inducing DNA transfer or recombination events, and providing virulence attributes (Wendling et al.; Canchaya et al. 2003; Banks et al. 2003; Eggers et al. 2016; Dedrick et al. 2017). For example, the pathogenicity of some strains of *Staphylococcus aureus* is reliant on the presence of integrated prophage sequences (Bae et al. 2006). In the active stage, the result of phage lysis significantly impacts microbial communities by turning over essential nutrients, especially carbon, nitrogen and sulfur (Emerson et al. 2018; Waldbauer et al. 2019; Anantharaman et al. 2014; Roux et al. 2016; Trubl et al. 2018; Suttle 2007). Lysis of bacterial populations likewise alters whole microbiomes by diversifying community structures and expanding niche opportunities (Gobler et al. 1997; Zimmerman et al. 2020). For example, the “Kill the Winner” model of virus population growth suggests that dominant bacterial populations are more susceptible to phage predation, which will facilitate expansions of less abundant taxa as the dominant populations are lysed (Thingstad and Lignell 1997; Thingstad 2000; Breitbart et al. 2018). Despite the importance of phage lysis on microbial communities, the proportion of lysis by prophages entering the lytic cycle is unclear. As opposed to strictly lytic phages, it remains difficult to associate prophages with active lysis. This is because prophage genome abundance can fluctuate according to host genome replication in the absence of lysis, whereas lytic phages, with few exceptions, must lyse a host in order to increase the abundance of their genomes.

In addition to traditional approaches such as isolation of phages, advances in high throughput metagenomic sequencing have sped up the ability to identify a large diversity of lytic and lysogenic phage sequences. Recently developed software have allowed for accurate characterization of prophages in both isolate and metagenomic assembled genomes, namely VIBRANT (Kieft et al. 2020a), VirSorter (Roux et al. 2015), PHASTER (Arndt et al. 2016) and Prophage Hunter (Song et al. 2019). Thus far these software have had the ability to extensively study the potential effects of prophages and begin to estimate the total diversity of prophages in nature. However, identifying the genome sequences of prophages does not provide context to their *in situ* state of being in the lysogenic or lytic stage of infection. This information is vital as it distinguishes which prophage or phage populations are actively impacting a microbial community through lysis events. Moreover, with the exception of Prophage Hunter, current software cannot distinguish prophage genomes that have become “cryptic”, or those that have lost functional abilities to enter the lytic stage (Noguchi and Katayama 2016; Ragunathan and Vanderpool 2019; Wang et al. 2010). Yet, Prophage Hunter still cannot identify if a given prophage is active, only if it may have the ability to do so.

Providing context to the infection stage of a prophage is imperative for accurate conclusions on its role in effecting its host and the microbial community. For example, identifying a prophage encoding a virulence factor or metabolic gene may have important implications for its role in manipulating its host’s pathogenic interactions, metabolic transformations, and impacts on nutrient and biogeochemical cycling. In order to place the prophage into context within the microbial community it would be necessary to first determine which stage the prophage is in, namely lytic or lysogenic. Assuming that all identified prophages are in a lytic stage could lead to misrepresentations or misinterpretations of the data if the prophage is actually dormant, or even cryptic.

Here, we present the software PropagAtE (Prophage Activity Estimator). PropagAtE uses genomic coordinates of integrated prophage sequences and short sequencing reads to estimate if a given prophage was in the lysogenic (dormant) or lytic (active) stage of infection. PropagAtE was designed for use with metagenomic data but can also use other forms of genomic data (e.g., sequence data from isolated microorganisms). When tested on systems with known active prophages PropagAtE was fully accurate in determining prophages that were active versus dormant, regardless of read coverage depth. No active prophages were identified in control systems encoding prophages that were known to be dormant. PropagAtE was also utilized to identify active prophages in several metagenomes, including the adult and infant human gut, murine gut and three different peatland soil environments. We show that specific prophages can be identified across various environments, and that activity of those prophages were restricted to particular environment types. Finally, we show that identifying the retention of a prophage over time does not necessarily indicate activity over time. PropagAtE is freely available at https://github.com/AnantharamanLab/PropagAtE.

## RESULTS

### Conceptualization of PropagAtE

Temperate phages that are integrated exist as a component of their host’s genome. When the host genome replicates, the prophage is also replicated likewise in a one-to-one ratio. As a result, when sequencing the host genome the prophage region and the flanking host region(s) are represented equally. Upon activation and entry into the lytic cycle, the prophage sequence is independently replicated for phage propagation and assembly into new virions. At this stage within the host cell there will be one host genome equivalent for multiple phage genomes regardless of whether lysis has occurred yet or not. Following lysis, virions containing phage genomes are released into the surrounding environment. These released genomes continue to represent the ratio of prophage to host genome copies if these prophage genomes are still included in the metagenome (**Fig. 1A**).

**Figure 1.**
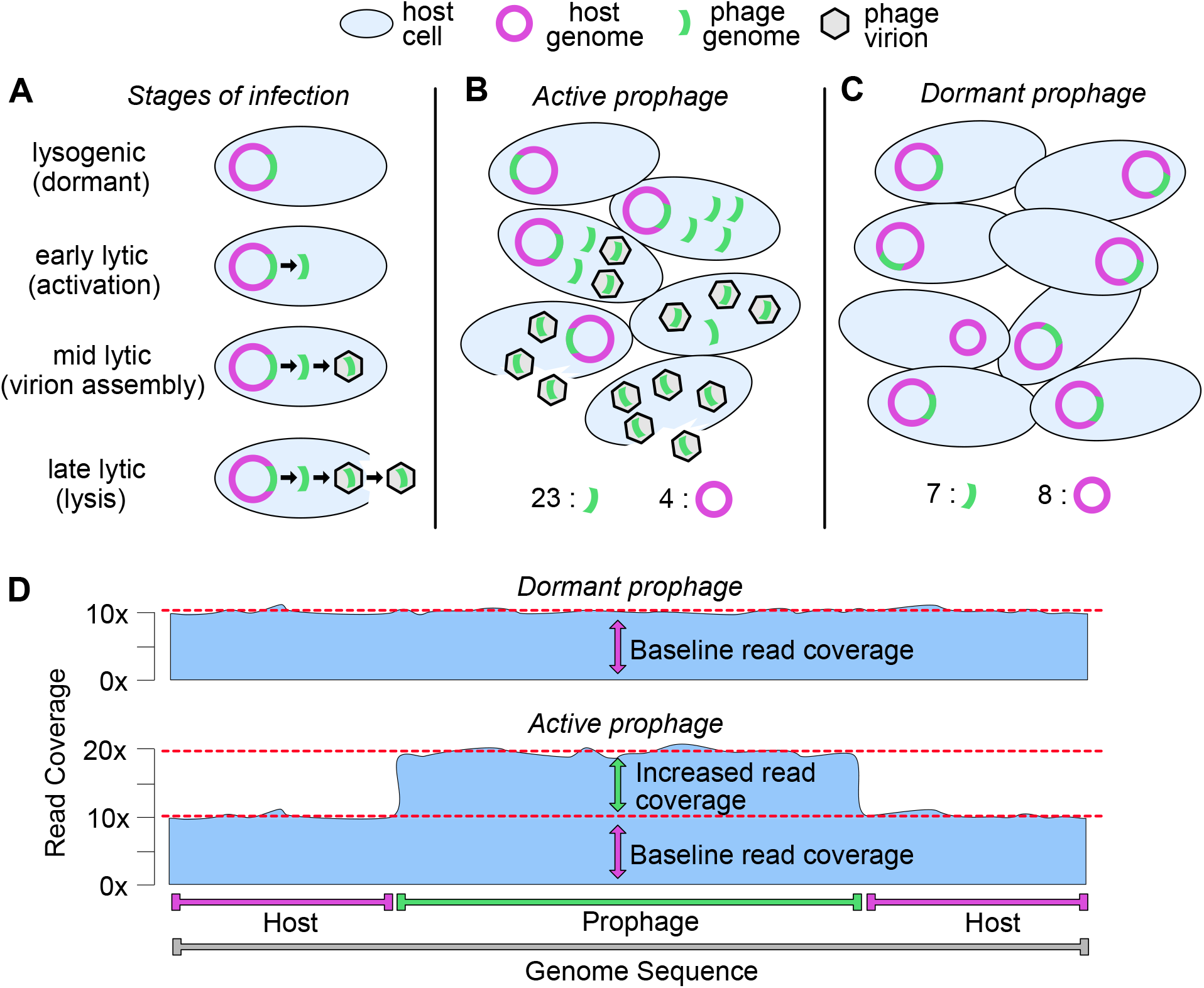
Conceptualization of PropagAtE mechanism. (A) Stages of integrated prophage infection from the lysogenic (dormant) to lytic (active) stages. Over the course of infection the prophage:host genome copy ratio increases. (B) Microbial community structure with an active prophage, from phage activation to lysis. The prophage:host genome copy ratio increases to greater than 1:1 through phage genome replication and host genome degradation. (C) Microbial community structure with a dormant prophage in which the prophage:host genome copy ratio is near 1:1. Here, one host is depicted as having cured the prophage from its genome.

The specific ratio of phage to host genomes depends on many factors. One major factor is the burst size of a given phage, or the number of virions released from a lysed host. Phage burst sizes can range from fewer than ten in the case of crAssphage that infects *Bacteroides intestinalis* (Shkoporov et al. 2018), to many thousands in the case of phage MS2 that infects *Escherichia coli* (Jenkins et al. 1974). Another factor, utilized by many phages including those that infect marine cyanobacteria, is that the host genome is degraded during the lytic stage to supply nucleotides to the replicating phage genomes which will further increase the prophage to host genome copy ratio (Weed and Cohen 1951; Zborowsky and Lindell 2019). Thus, during the lytic stage of phage propagation as well as post-lysis, the ratio of prophage to host genome copies will become skewed in favor of prophage genomes (Waller et al. 2014; Hertel et al. 2015). This will lead to a prophage:host genome copy ratio significantly greater than 1:1 (**Fig. 1B**). If the prophage was in a dormant stage of infection the prophage:host genome copy ratio would be approximately 1:1 (**Fig. 1C**). This is likewise dependent on various factors, such as the ability of some members of the host population to “cure” (i.e., remove) the prophage from its genome. Despite nuances in specific prophage:host genome copy ratios, active prophages will yield a ratio greater than 1:1 whereas dormant prophages will yield a ratio near 1:1.

Whether or not the prophage:host genome copy ratio is skewed can be identified using statistical analyses of aligned sequencing read coverage after genome sequencing and read alignment. After sequencing and assembly of a system (e.g., isolated bacteria culture, complex microbiome, etc.), the integrated prophage sequence will assemble as a component of the host genome in a ~1:1 ratio, regardless of activity. However, if a prophage has activated then the resulting phage genome copies contained in virions are identical to the integrated prophage sequence. Therefore, read alignment to the assembly will recruit reads to the prophage and host regions in a ratio indicative of the stage of infection. During the lysogenic stage where the prophage is dormant, read recruitment will generate even coverage across the regions. Conversely, a prophage that has entered the lytic, active stage will generate an uneven read recruitment skewed towards greater coverage at the prophage region only (**Fig. 1D**). Read alignment will not determine the true prophage:host abundance, but it can quantify a relative ratio to accurately determine stage of infection.

### Overview of PropagAtE’s workflow

Differentiating active prophages from those that are dormant is essential for accurate representation and evaluation of individual-cell and community-level systems. PropagAtE provides the first automated platform for the identification of active prophages that is scalable for isolate genomes or complex metagenomes. Since most prophages exist as an integrated (i.e., connected) element of a host genome, the read coverage from the prophage and host sections can be compared in a one-to-one manner to estimate a genome copy ratio. PropagAtE utilizes the ratio of prophage:host read coverage along with the ratio’s effect size (i.e., significance of the ratio) to designate if a given prophage was dormant or active. The PropagAtE workflow can be simplified into three general steps: data input, read alignment processing, and results output (**Fig. 2A**). Users are given two options for data input: (1) genomes/scaffolds of host sequences with raw short sequencing reads or (2) a pre-generated alignment file in SAM or BAM format. If given the former input, reads will be aligned using Bowtie2 (Langmead and Salzberg 2012) to generate a SAM file. If given an input in BAM format it will first be converted to SAM before proceeding (Li et al. 2009).

**Figure 2.**
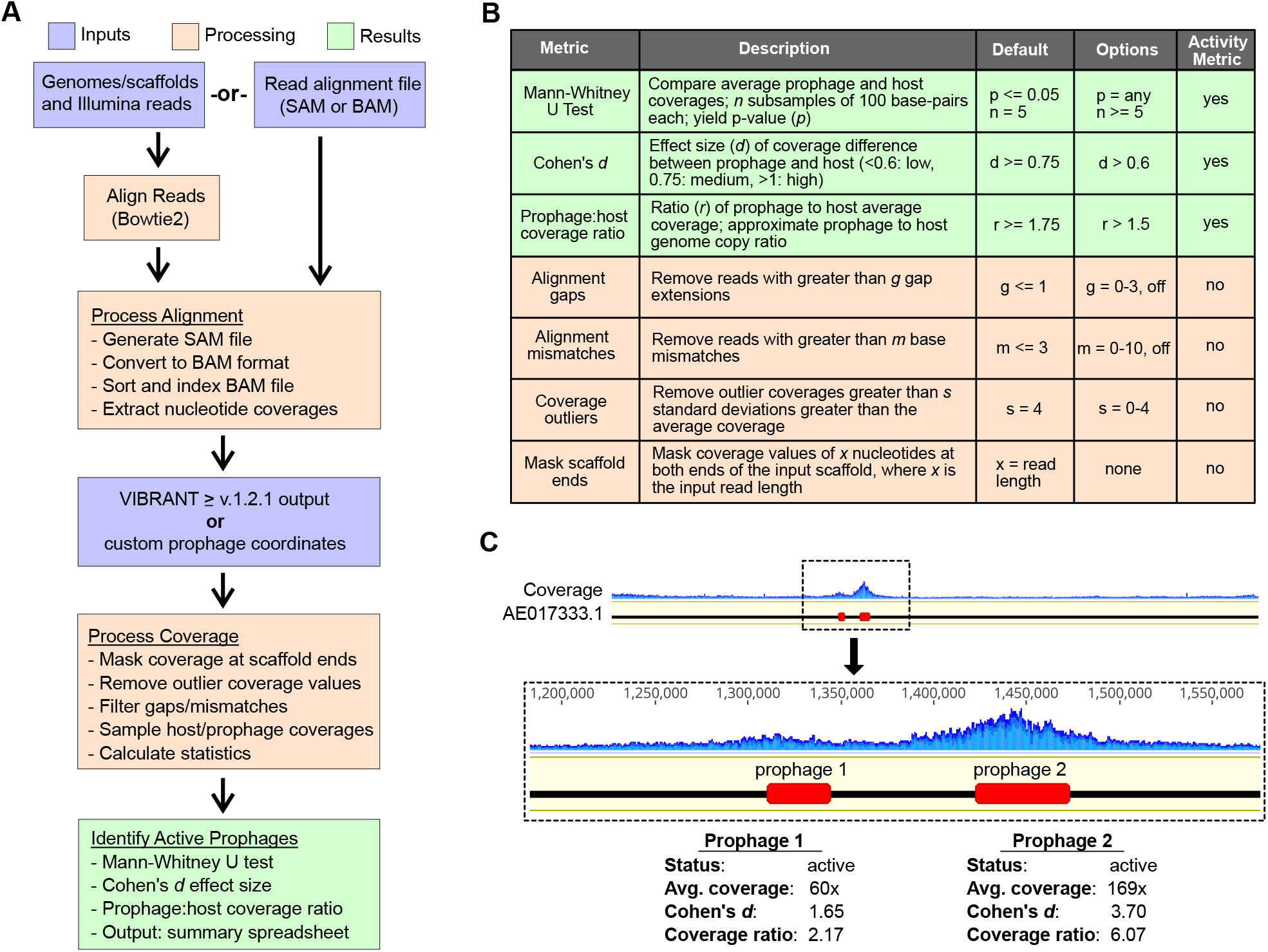
Workflow and implementation of PropagAtE. (A) Workflow of PropagAtE including data input, read alignment processing and results output. (B) Summary table of PropagAtE activity metrics, processing metrics, thresholds and options. (C) Example of read coverage profiles for two active *Bacillus licheniformis* DSM13 prophages, respective to the conceptual diagram in **Figure 1D**.

Using the SAM file, either generated or supplied by the user, aligned reads exceeding the gap and/or mismatch thresholds are removed. Following gap/mismatch filtering, the alignment is converted to BAM format and coverage per nucleotide is extracted including all nucleotides with zero coverage. To eliminate noise, coverage values at the sequence ends are trimmed off. The number of coverage values trimmed per end is equal to the input read length. Then, users are given two options for prophage coordinate data input: (1) direct results from a VIBRANT (v1.2.1 or greater) analysis (Kieft et al. 2020a) or (2) a manually generated coordinate file of a specified format. In cases for which multiple prophages are present on a single genome/scaffold, all prophage regions are considered independently. In addition, the host region is segmented to exclude all prophage regions, but each segment is considered as a single, cohesive host sequence. That is, if two or more prophages are present on a single host scaffold, neither prophage will interfere with the other in terms of coverage value calculations and each prophage is compared to an identical prophage-excluded host region.

Next, outlier coverage values are removed from consideration before calculating statistical metrics. For each prophage and host pair, various metrics are calculated including average coverages, median coverages, coverage standard deviations and prophage:host coverage ratio. Each prophage’s activity is estimated according to (1) the prophage:host coverage ratio, (2) Cohen’s *d* effect size of the coverage difference and (3) the combined Mann-Whitney test *p*-value. Along with the prophage:host coverage ratio, the effect size of the difference between the prophage and host coverages is compared using Cohen’s *d* metric. For Mann-Whitney statistical analysis, at least five random samples of 100 coverage values are taken from both the prophage and host sequence regions and the resulting *p*-values per subsample are combined. Prophages exceeding the default or user-set thresholds for all three metrics are considered active (**Fig. 2B**).

### Read alignment can visualize active prophages

Two activated prophages in the genome of *Bacillus licheniformis* DSM13 were used to visualize active prophage identification using PropagAtE (**Fig. 2C**). Visualization of the read coverage at each nucleotide in the genome clearly depicted coverage spikes exclusively at the prophage regions. The example prophages existed in close proximity to each other and had differing average coverages (60x and 169x). Both example prophages likewise met the minimum prophage:host coverage ratio (2.17 and 6.07) and Cohen’s *d* effect size (1.65 and 3.70) thresholds. These results are in line with the conceptualization of the workflow seen in **Fig. 1D**.

### Positive control tests for prophages from isolate genomes

Positive control tests were utilized in order to set threshold boundaries for PropagAtE to identify active prophages as well as assess PropagAtE’s recall rate. Positive control samples were considered as those for which DNA from both an active prophage and its host were extracted and sequenced in tandem. This method best represents metagenomic samples in which all DNA is extracted and sequenced together. In addition, extraction of both host and free phage DNA together is essential for positive tests because this method will best depict the most accurate prophage:host ratio. Three model systems for which sequencing data was publicly available were identified for use as positive controls. All experiments and sequencing were performed elsewhere (Hertel et al. 2015; Gutiérrez et al. 2018; Ho et al. 2016) (**Supplemental Table S1**). Each system, since they represent isolate bacteria, have a much higher read coverage compared to a typical metagenome assembled genome. To ensure validation of PropagAtE for both isolate and metagenomic samples, two tests per system were done. One was done with all available reads (“full reads”) and another was done with a random subset of 10% of the reads (“10% reads”). All PropagAtE results for positive control tests can be found in **Supplemental Table S2**.

The first system we tested was *Bartonella krasnovii* OE1-1 and its prophage (Gutiérrez et al. 2018). In triplicate, the bacteria were either induced for prophage using mitomycin C or uninduced as controls. For the induced prophages, the prophage:host coverage ratios were relatively even between the three samples (1.78x, 1.83x and 2.03x). Likewise, in the uninduced control samples the prophage:host coverage ratios depicted nearly equal coverage (1.08 and 1.03) except one sample with a low ratio (0.43) (**Figs. 3A, B**). This suggests the method is reliable across multiple samples or time points for the same phage. The ratio effect size, using Cohen’s *d* metric, indicated that the prophage:host coverage ratios observed were significant in their difference. For the induced prophages the effect sizes were greater than one (1.13, 1.09, 1.14) indicating a high dissimilarity between the prophage and host coverages. The uninduced controls’ effect sizes were low (0.41, 0.17 and 0) (**Fig. 3B**). When 10% of the reads were randomly sampled for PropagAtE, the induced (1.75x, 1.84x and 2.05x) and uninduced (1.04x, 1.08x and 0.45x) results were essentially equivalent to that of the full read set in terms of prophage:host coverage ratios (**Figs. 3C, D**) and only marginally lower effect sizes (induced: 0.99, 0.99 and 1.03; uninduced: 0.11, 0.21 and 0) (**Fig. D**). This further indicates that high read coverage is not essential, nor significantly impacts, the outcome of analysis.

**Figure 3.**
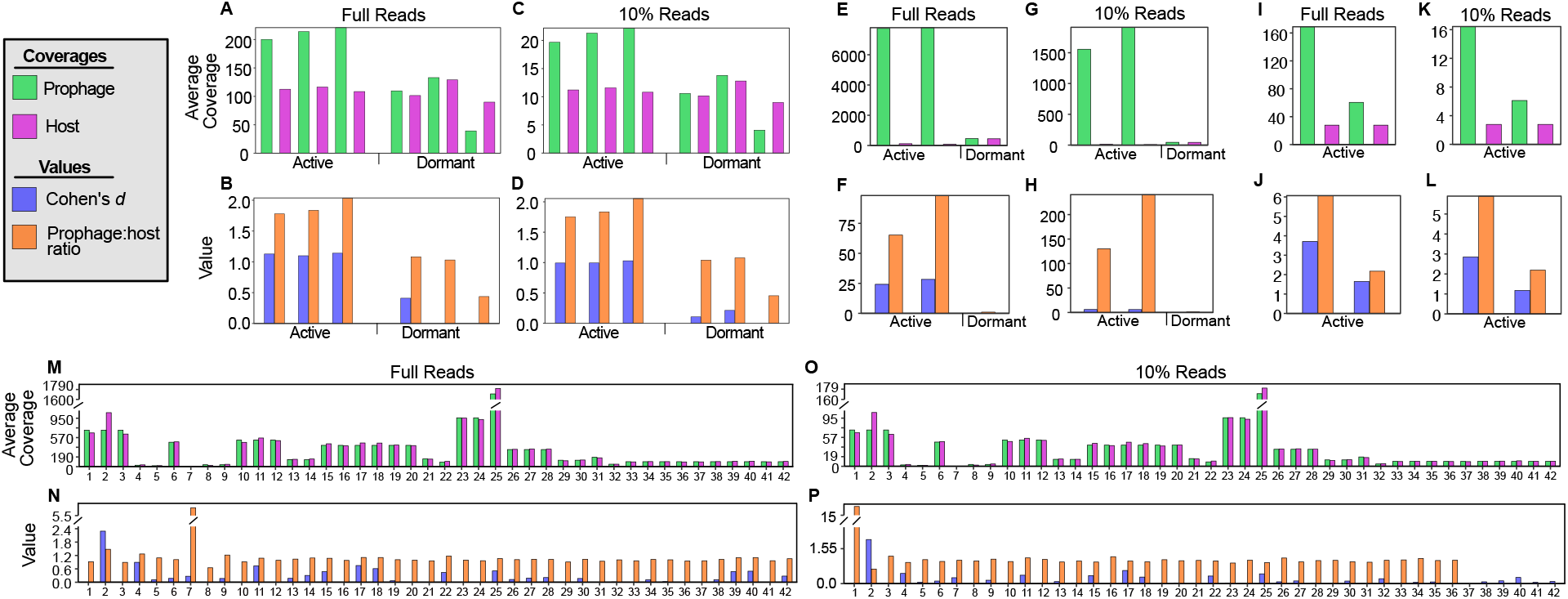
Positive and negative control results. (A-L) All positive control and (M-P) all negative control test results. For both prophages and hosts average read coverage is shown (green and purple bars) as well as Cohen’s *d* effect sizes and prophage:host coverage ratios (blue and orange). For the positive control tests, samples are labeled as containing active or dormant prophages.

The second system we tested was *Lactococcus lactis* MG1363 and its prophages (Ho et al. 2016). Similar to the previous system, in one sample the prophage was induced with mitomycin C and another was used as an uninduced control. The induction sample was sequenced 1- and 2-hours post-induction for a total of two positive samples. For the induced samples the resulting prophage:host coverage ratios were high and increased over time (65x and 98x). In the uninduced control the prophage:host coverage ratio was, as seen with the previous system, nearly equal (1.02) (**Figs. 3E, F**). The effect size of the ratio for the induced samples were high and increased over time (24.2 and 28.3) while the effect size of the control sample ratio was low (0.10). The results from 10% subsampled reads yielded slightly different but equally determinant values. The prophage:host coverage ratios for the induced samples increased (130x and 241x) while the control ratio remained constant (1.02) (**Fig. 3G, H**). The effect sizes for the induced coverage ratios were smaller (5.96 and 5.92) but were nonetheless very high, while the control’s effect size remained the same (0.10) (**Fig. 3H**).

The third system we tested was *Bacillus licheniformis* DSM13 and its prophages (Hertel et al. 2015). Here, both prophages were spontaneously activated at 26°C and no control was used for comparison. The prophage:host coverage ratios (2.17x and 6.07x) as well as the corresponding effect sizes (1.65 and 3.70) were significant (**Figs. 3I, J**). This was likewise observed when 10% subsampled reads were used (ratio: 2.19x and 5.90x; effect size: 1.18 and 2.85) (**Figs. 3K, L**). Although the available control sample size could not strictly designate a true discovery rate, all controls tested resulted in perfect accuracy and recall.

### Negative control tests for prophages from isolate genomes

Negative control tests were utilized in order to set threshold boundaries for PropagAtE to identify dormant prophages as well as assess PropagAtE’s specificity. Several negative control samples were used for testing in addition to the control samples presented above. Negative controls were considered as those in which a bacterial genome encoding at least one prophage was sequenced in the absence of known prophage induction. A total of 19 diverse bacterial genomes encoding 42 predicted prophages were used. As before, each system was tested with a set of all reads as well as smaller dataset containing 10% randomly subsampled reads. All sequencing was performed elsewhere (**Supplemental Table S1**). All PropagAtE results for negative control tests can be found in **Supplemental Table S2**.

When using the complete reads sets, all 42 prophages were found to be dormant. Average prophage (1810x to 0.80x) and host (1705x to 0.13x) coverages ranged considerably (**Fig. 3M**). All prophage:host coverage ratios were below 1.5 except for one, which was 5.80. However, the effect size of the high prophage:host coverage ratio was only 0.27. All coverage ratio effect sizes ranged from 2.29 to 0 (**Fig. 3N**). A total of four prophages had effect sizes greater than the threshold, but actual prophage:host coverage ratios that were too small (1.47x to 1.08x). For the 10% subsampled read results, the prophage:host coverages ranged from 1.48x to 0.65x, with the same exception as before (15.4x) (**Fig. 3O**). Again, the effect size of the latter ratio was 0.26. The remaining coverage ratio effect sizes ranged from 1.95 to 0 (**Fig. 3P**). Given that all prophages were identified as dormant these results suggest that the two metrics, prophage:host coverage ratio and corresponding effect size, function adequately in a check and balance system with each other. Prophages with significantly high prophage:host coverage ratios had insignificant effect sizes, and vice versa. Likewise to the positive control tests, the observed false discovery rate was zero, though the true accuracy of PropagAtE is likely small but greater than zero.

### Applying PropagAtE to identify active prophages in metagenomes

PropagAtE was designed to rapidly assess the activity of prophages in metagenomes in a high-throughput manner. Additionally, PropagAtE can also identify active prophages in genomes of cultivated organisms, irrespective of the manner of prophage induction (i.e., spontaneously or experimentally induced). To validate the broad utility of PropagAtE, we demonstrate its application on 348 metagenomic samples from a variety of environments: adult and infant human gut, murine gut, and peatland soil (Emerson et al. 2018; Kim and Bae 2018; He et al. 2017; Gasparrini et al. 2019; Woodcroft et al. 2018; Feng et al. 2015) (**Table 1, Supplemental Table 1**). A total of 377 semi-redundant prophages were identified as active across all samples. Per sample, the percent of prophages that were active ranged from 0% to 22% with an average of 2.2% (**Fig. 4**). The murine gut had the most active prophages per sample with an average of 12.2% whereas all human gut samples had a combined average of 1.5%. These results show that for metagenomic samples most prophages identified as integrated into a host genome are dormant. All PropagAtE results for metagenomic samples can be found in **Supplemental Table S3**.

**Figure 4.**
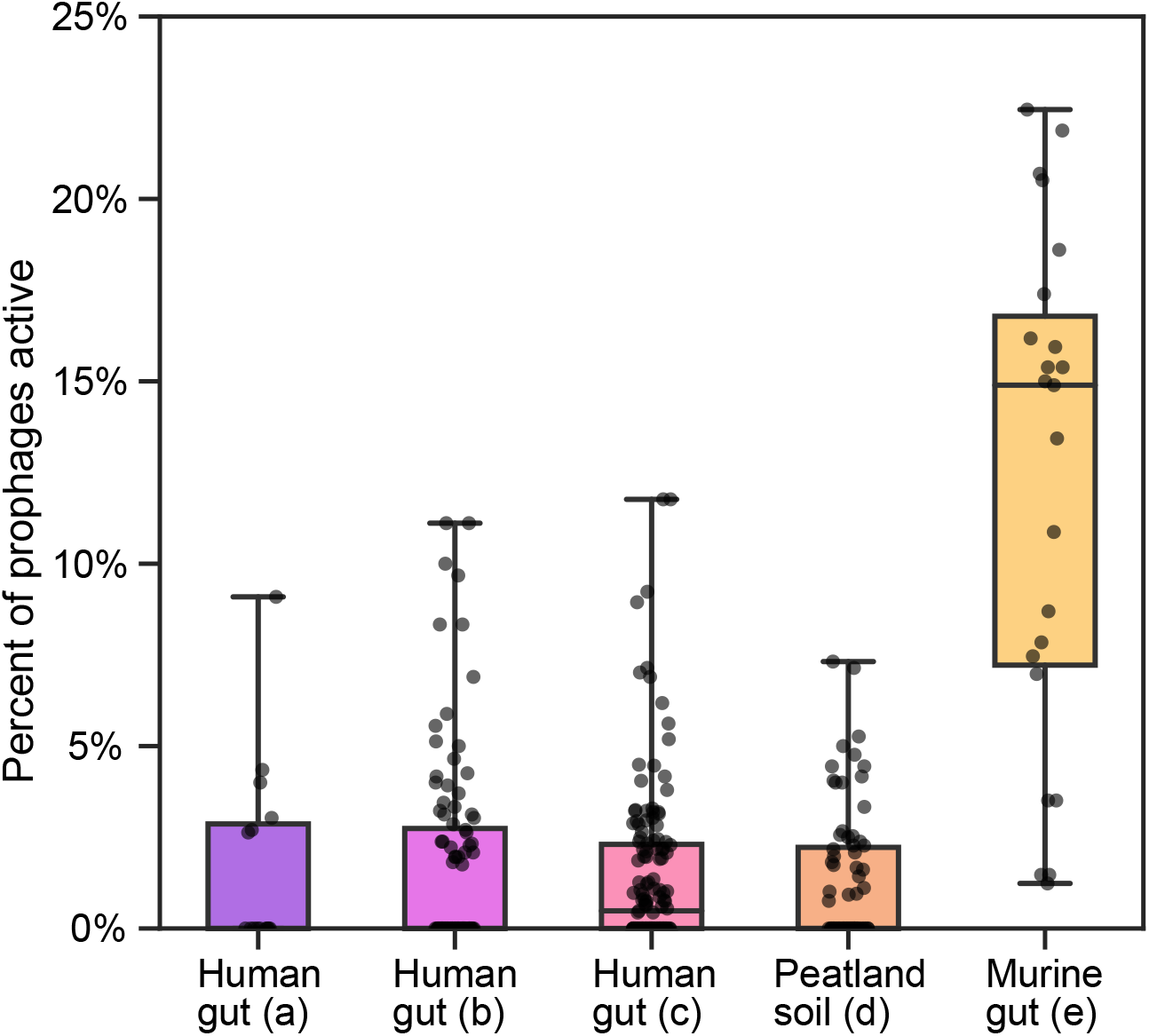
Percent of prophages identified as active in metagenomic samples. Five sets of metagenomic samples are compared with each dot representing a single sample. Identifier labels a-e on the x-axis correspond to the final column in **Table 1**.

**Table 1.** Summary of metagenomic sample datasets. The environment type, description of the dataset and total number of samples per metagenomic dataset are provided. The final column, except for (f), corresponds to labeling in **Figure 4**.

| Environment | Description | No. of samples | Citation | Label |
| --- | --- | --- | --- | --- |
| Human gut (fecal) | Adult individuals with colorectal adenoma, carcinoma or healthy controls | 15 | (Feng et al. 2015) | a |
| Human gut (fecal) | Adult individuals with Crohn's Disease or healthy controls | 96 | (He et al. 2017) | b |
| Human gut (fecal) | Infant individuals given antibiotics or untreated controls | 139 | (Gasparini et al. 2019) | c |
| Peatland (soil core) | Soil cores of bog, fen and palsa environments | 75 | (Woodcroft et al. 2018; Emerson et al. 2018) | d |
| Murine gut (fecal) | Microbial and virome fractions of murine gut | 23 | (Kim and Bae 2018) | e |
| Human gut (fecal) | Time series of adult individuals with Crohn's Disease | 12 | (Ijaz et al. 2017) | f |

For metagenome datasets with various conditions (e.g., peatland palsa, bog and fen habitats) no significant difference was observed in the total number of active prophages per condition (**Supplemental Fig. S1, Supplemental Table S4**). However, utilizing PropagAtE to identify which sets of prophages are active yielded interesting results. For example, differences in specific populations of prophages between peatland soil bog, fen and palsa habitats was observed. Between these habitats, three major populations of active prophages were identified according to the host bacterial species. Note that accurate host identification can be made since the prophages are integrated into the host genome. These three populations infected Myxococcales, Acetobacteraceae and Acidimicrobiaceae. Active Myxococcales prophages were found exclusively in fen habitats. Active Acetobacteraceae prophages were found predominately in palsa habitats and somewhat in bog, whereas active Acidimicrobiaceae prophages were found predominately in bog habitats and somewhat in palsa. The latter two were not identified in fen habitats (**Fig. 5, Supplemental Table S5**). Notably, 167 Myxococcales, 322 Acetobacteraceae and 429 Acidimicrobiaceae dormant prophages with average coverages greater than zero were identified. This indicates that although a given prophage population may be found across multiple samples they may only be active in specific habitats.

**Figure 5.**
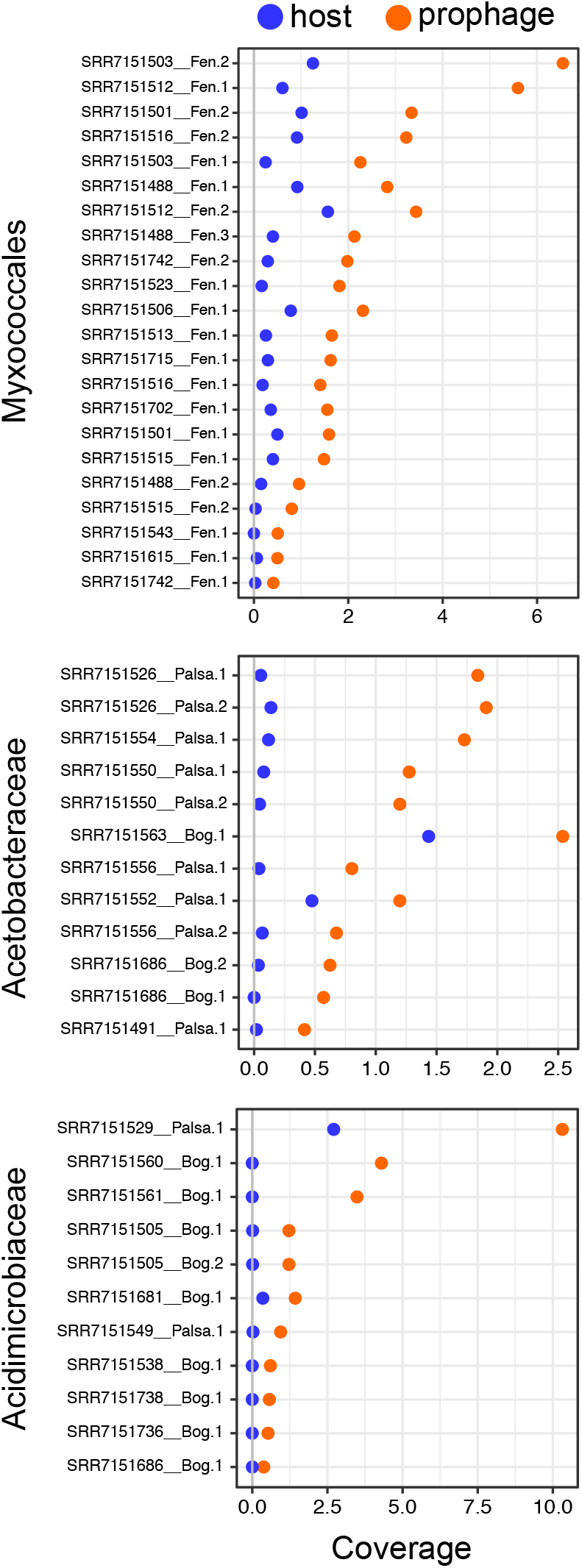
Active prophages identified in peatland soil habitats. Active prophages integrated in Myxococcales, Acetobacteraceae and Acidimicrobiaceae genomes are shown. Each pair of dots represents a single prophage population, with blue dots (left) depicting host coverage and orange dots (right) depicting prophage coverage.

### Estimating prophage activity over time

To further explore the activity of specific prophage populations over time, a sixth set of metagenomic samples was used (Ijaz et al. 2017). This set included human gut fecal samples from three different children with Crohn’s Disease. For each individual, four time series samples were taken at approximately days 0, 16, 32 and 54. Among all three individuals, a total of 11 unique prophages were identified across all four time points. None of the 11 prophages were shared between two or more individuals. Therefore, these 11 prophages were found to be consistently present and retained stably over time. All prophage populations encoded hallmark phage proteins, nucleotide replication proteins and lysis proteins, indicating they likely have the ability to activate (i.e., not cryptic). For most populations, genes for integration were also identified (**Supplemental Fig. S2A, Supplemental Table S6**). Furthermore, one prophage population encoded the auxiliary metabolic gene *cysH* for assimilatory sulfate reduction, a metabolic process that can yield hydrogen sulfide, which has been implicated in exacerbating inflammatory bowel diseases such as Crohn’s Disease (Guo et al. 2016; Wallace et al. 2009). Another prophage population encoded a RhuM family virulence protein. Genome alignment of each prophage population yielded 99.8-100% identity with a maximum number of two nucleotide differences between members of a population (**Supplemental Fig. S2B**). However, PropagAtE identified none of these prophages to be active at any time point. This conclusion is important as it suggests that the prophages, in addition to the *cysH* and *rhuM*-like genes, were present but may not have been actively impacting the microbial community. The lack of sequence diversification likewise suggests the prophage populations were primarily dormant over time since active phage genome replication typically results in nucleotides changes. However, the minor nucleotide differences may have resulted from alignment or sequencing error, or from prophage activity between the time points sampled.

### Sequencing depth does not correlate with total active prophages

As a final validation test, we examined if the total number of sequencing reads (i.e., sequencing depth) had an impact on the total number of active prophages identified. It may be assumed that since PropagAtE relies on read coverage, samples with a greater number of reads would identify disproportionately more active prophages. Using the five sets of metagenomic samples (**Table 1**) we correlated the total number of reads used by PropagAtE to the total number of active prophages identified. Four of the five sets of metagenomic samples yielded near linear, flat trends indicating no correlation between total reads and total active prophages. The fifth set, representing infant gut samples, depicted more of a trend towards a correlation between more reads and more active prophages. However, the trend was not significant (**Supplemental Fig. S3, Supplemental Table S7**).

### PropagAtE run time

Efficiency and quick run speed are essential for large-scale metagenomic workflows. PropagAtE was designed to meet the needs of these analyses, such as those with many samples or large file sizes. PropagAtE is likewise scalable for smaller datasets. To show this we estimated the total run time for various isolate and metagenome samples. For isolate samples, run time typically was between 5 and 10 minutes with reads as the input, or about 1 minute with an alignment format file (i.e., BAM format) as the input. For metagenomes, the run time was slightly longer at 10-15 minutes (**Supplemental Table S8**). The main factor affecting run time is read alignment performed by Bowtie2. It is important to note that the run time for large read dataset inputs significantly improves when utilizing the multi-threading feature, though alignment format file inputs are not affected. Without multithreading, large read dataset inputs can result in long run times.

## DISCUSSION

Phages are key contributors to microbiome dynamics in essentially all environments on Earth (Suttle 2007; Wilhelm and Suttle 1999; Trubl et al. 2018; Roux et al. 2016; Paez-Espino et al. 2016; Shkoporov et al. 2019; Kieft et al. 2021, 2020b). With the availability of high-throughput sequencing and newly developed software tools we have the ability to identify and study these diverse phages (Arndt et al. 2016; Kieft et al. 2020a; Roux et al. 2015; Song et al. 2019). This includes both strictly lytic phages as well as integrated prophages. However, little emphasis has been placed on identifying which populations of identified prophages are actively replicating as opposed to existing in a dormant or cryptic stage of infection.

Here we have presented the software tool PropagAtE for the estimation of activity of integrated prophages using statistical analyses of read coverage. Although the concept of using read coverage to predict prophage activity is not new (Hertel et al. 2015), PropagAtE is the first benchmarked implementation of the method into an automated software for use with large datasets, such as metagenomes. PropagAtE functions by quantifying the relative genome copy ratio between a prophage region compared to a corresponding host region. Only prophages that have activated and begun propagation (e.g., genome replication and virion assembly) will yield prophage:host ratios sufficiently greater than 1:1. The prophage:host genome copy ratio, estimated by using read coverage ratios, as well as the ratio’s effect size are used to classify a prophage as active or dormant. We provide evidence to show that PropagAtE is fast, sensitive and accurate in predicting prophages as active versus dormant and have applied the method to various metagenome samples.

Identifying which prophage sequences are active versus dormant in a sample provides several benefits. Namely, assuming that all identified prophages are active is an overestimation and will lead to a misrepresentation of the *in situ* dynamics of a microbial community. For example, we show here that Myxococcales prophages identified in fen, bog and palsa soils are strictly active in fen habitats. Despite their sequence identification in bog and palsa samples, the most accurate representation of the prophages is to conclude that their effect on the resident microbial communities is likely restricted to fen samples where host lysis is occurring. We likewise show this for 11 unique prophages that were found to be dormant across multiple sampling time points in human gut environments. Another benefit includes making accurate conclusions on the role of host bacteria in a given sample. Foremost, prophages can be responsible for the virulence of multiple human pathogens, such as *Clostridioides difficile, Clostridium botulinum, Staphylococcus aureus* and *Corynebacterium diphtheria* (Schroven et al. 2021; Lindsay et al. 1998; Freeman 1951; Brüssow et al. 2004; Sakaguchi et al. 2005; Riedel et al. 2017). Although some virulence effects are present during prophage dormancy and expression of specific genes, many require activation of the prophage. In addition to virulence, bacteria actively infected by a phage can have a modified metabolic landscape compared to bacteria uninfected or harboring a dormant prophage. Several examples include the phage-directed regulation of sulfur, carbon, nitrogen and phosphorus metabolism in various cyanobacteria and enterobacteria (Waldbauer et al. 2019; Howard-Varona et al. 2020; Thompson et al. 2011; Stent and Maaløe 1953). This distinction is vital when assessing the role of the microbial community in an environment. Related to this, activity can provide context to any auxiliary metabolic genes identified on the prophage genome, such as *cysH* for assimilatory sulfate reduction described here. In the human gut specifically, identifying phage-encoded genes for sulfur metabolism may have important implications for the health of the gastrointestinal tract and a phage’s role in the manifestation or perturbation of diseases (Clooney et al. 2019; Kieft et al. 2021). If a prophage encoding an auxiliary metabolic gene is identified, determining the stage of infection of the prophage can provide context to the effect of the auxiliary metabolic gene.

It is important to point out several unavoidable caveats to the implementation of PropagAtE. First, accurate prophage:host genome copy ratio estimations are inhibited if the sample is size fractionated before sequencing. For example, many aquatic samples are size fractionated by filtering onto a 0.2-micron filter. In these cases, only pre-lysis infections will be picked up by read coverage because the genomic content present in released virions will likely pass through the 0.2-micron filter. Second, not all prophages exist as integrated sequences, such as those that are episomal. Prophages that are episomal do not have attached host sequence and therefore cannot have prophage:host read coverage compared in a one-to-one manner, and for metagenomes cannot have accurate host prediction. This also applies to prophages that do not assemble as integrated components of a host scaffold. However, it is worth noting that for integrated prophages PropagAtE functions whether the host region flanks the prophage on one or both sides. Third, though not verified, is that inactive prophages may be more likely to assemble with a host scaffold. Since active prophages lyse their host and potentially degrade their host’s genome, more activity of a prophage may lead to a lower probability of assembling as an integrated prophage. Due to the caveats presented, PropagAtE is intended to be used for identifying active prophage sequences rather than assessing the total number or fraction of prophages that are active in a sample. In this context PropagAtE performs with little to no observed error. Finally, PropagAtE has been developed and tested using short read sequencing data and is not yet suitable for long read analyses.

Overall, our results demonstrate that PropagAtE will facilitate the accurate characterization and study of viruses in microbiomes and nature. Examples of future applications of PropagAtE include the exploration of prophages in human health and disease, detection of environmental and chemical triggers for induction of prophages, phage therapy research (for disqualifying prophages), and in environmental systems research.

## METHODS

### Datasets used for control tests

All datasets, genomes and reads, used for positive and negative control tests were acquired from publicly available datasets on NCBI databases (O’Leary et al. 2016; Clark et al. 2016). See **Supplemental Table S1** for details of studies and accession numbers. VIBRANT (v1.2.1) (Kieft et al. 2020a) was used for identification and annotation of all prophages for both control datasets as well as metagenomes.

### Dependencies and equations

Bowtie2 (v2.3.4.1 in this study) (Langmead and Salzberg 2012) was used for read alignment. Samtools (≥ v1.11) (Li et al. 2009) was used for manipulation, conversion and reading of SAM and BAM alignment files. SciPy (Virtanen et al. 2020) was used for Mann-Whitney tests and other statistical analyses. Cohen’s *d* metric is used to calculate the effect size of prophage:host coverage ratios. Cohen’s *d* (Cohen 2013) is calculated using the following equation where 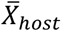 and 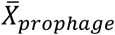 are the average read coverages of the host and prophage regions, and *S_host_* and *S_prophage_* are the standard deviations of the coverages:

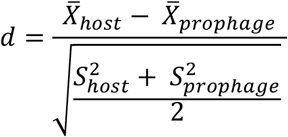

### Metagenome assembly and analyses

Metagenomes for the murine gut microbial fraction samples were assembled in this study. Details of raw read sets from murine gut samples used for assembly can be found in **Supplemental Table S1**. SPAdes (v3.12.0) (Nurk et al. 2017) was used for genome assembly (−−meta-k 21,33,55) and the resulting best scaffold assemblies were retained. The human infant gut and peatland soil metagenomes were assembled previously in their respective studies (Emerson et al. 2018; Gasparrini et al. 2019; Woodcroft et al. 2018). Both human adult gut metagenomes were assembled by Pasolli et al. (Pasolli et al. 2019).

For the human gut time series samples integrated prophages were predicted using VIBRANT (v1.2.1). To check for integrated prophage sequences that were not assembled with a host scaffold, integrated prophages were compared to all identified phages using dRep (v2.6.2, dereplicate – ignoreGenomeQuality-sa 90-pa 90) (Olm et al. 2017). Identical, non-integrated phage sequences were considered as a part of the same prophage population. Genome alignments were performed using progressive Mauve (default settings) (Darling et al. 2010).

### Visualization

Geneious Prime 2020.1.2 was used for visualization of example read coverage values. R package ‘ggplot2’, Matplotlib and Seaborn were used for visualization of graphs (Michael Waskom et al. 2017; Hunter 2007).

### Setting default thresholds for PropagAtE

PropagAtE has several, variable settings and thresholds that can be set by the user: number of gap extensions per aligned read, number of mismatched base pairs per aligned read, standard deviation threshold for outlier coverage removal, *p*-value cutoff for Mann-Whitney U test and number of coverage subsamples taken for the test, minimum prophage:host coverage ratio, and minimum Cohen’s *d* effect size. The default settings and setting options can be found in **Figure 2C**.

The first three settings are used for read alignment processing: number of gap extensions, number of mismatched base pairs and standard deviation threshold. The default number of gap extensions per aligned read is one or fewer. This setting is meant to be sensitive for accurate read alignment while allowing for minor errors. Likewise, the number of mismatched base pairs per aligned read is set to three or fewer in order to be sensitive and accurate. To designate the default to three, total mismatches per read between 0-4 were tested (**Supplemental Fig. S4**). Outlier coverage values are removed in order to take into account alignment errors that may incorrectly stack reads. The default for outlier coverage removal is set to four standard deviations above the calculated average in order to only remove artificially high coverage values. A fourth, immutable metric is masking of coverage values at genome/scaffold ends. This setting is particularly important for metagenomic scaffolds that likely represent partial sequences. For this metric, the estimated length of the input reads is used to mask (i.e., not consider for calculation) the respective number of coverage values from each scaffold end in order to account for lower coverage values at partial scaffold ends.

The final three settings are used for determination of prophage activity and significance: Mann-Whitney U test *p*-value, prophage:host coverage ratio and Cohen’s *d* effect size. The default *p*-value cutoff for the Mann-Whitney test is set to the standard value of 0.05 but can be set to any value. This metric for designating a prophage as active or dormant is the least sensitive. The default setting takes five subsamples of 100 individual nucleotide coverage values to generate *p*-values in order to avoid artificially low *p*-values from high *n* values (i.e., thousands of nucleotides per input genome/scaffold). The most important threshold is the prophage:host coverage ratio, which is set to 1.75 by default. The default was selected to be as close to the minimum requirement for designating true active prophages as active in control tests while maintaining a significant gap from true dormant prophages in order to reduce false positive identifications. Finally, Cohen’s *d* effect size setting is set to 0.75 which falls in the general range of “medium” significance. This threshold is useful for contextualizing prophage:host coverage ratios, especially for high-coverage genomes/scaffolds. Again, the default was selected according to control tests for reducing false positive identifications.

## DATA ACCESS

The PropagAtE software and associated files are freely available at https://github.com/AnantharamanLab/PropagAtE. All isolate and metagenome genomic sequences and reads used in this study are publicly available; see **Supplemental Table S1** for details. Additional details of relevant data are available on request.

## ACKNOWLEDGEMENTS

We thank the University of Wisconsin—Office of the Vice Chancellor for Research and Graduate Education, University of Wisconsin—Department of Bacteriology, and University of Wisconsin—College of Agriculture and Life Sciences for their support. K.K. is supported by a Wisconsin Distinguished Graduate Fellowship Award from the University of Wisconsin-Madison. We also thank Z. Zhou, A. Adams and R. Salamzade for their helpful feedback and discussions.

