## Supplemental Figures S1-S4 for "Deciphering active prophages from metagenomes"

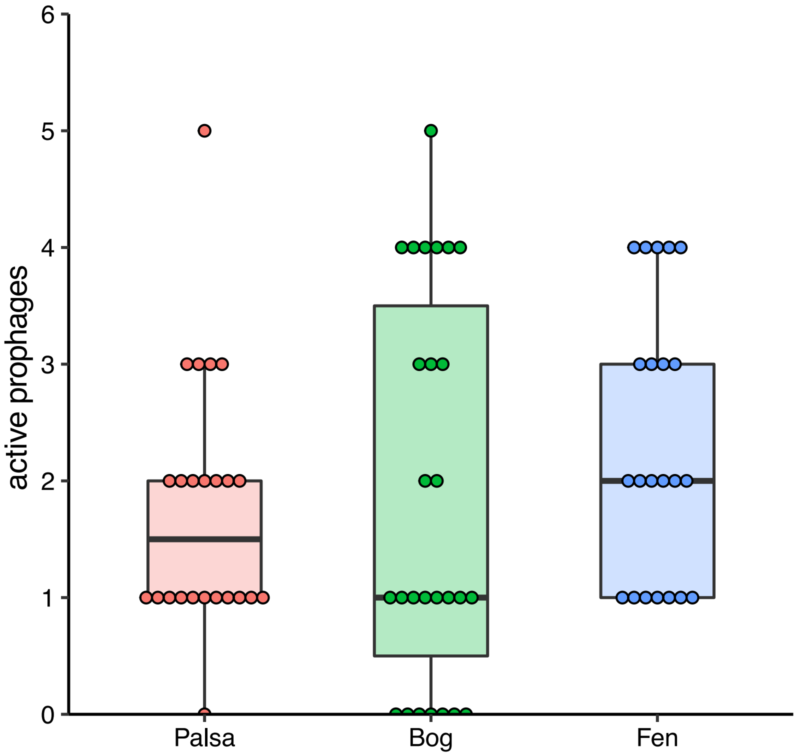


**Supplemental Figure S1. Total active prophages identified in peatland palsa, bog and fen habitats**. Each dot represents the total number of active prophages identified in a single metagenomic sample. No difference in total active prophages were identified between the environments.

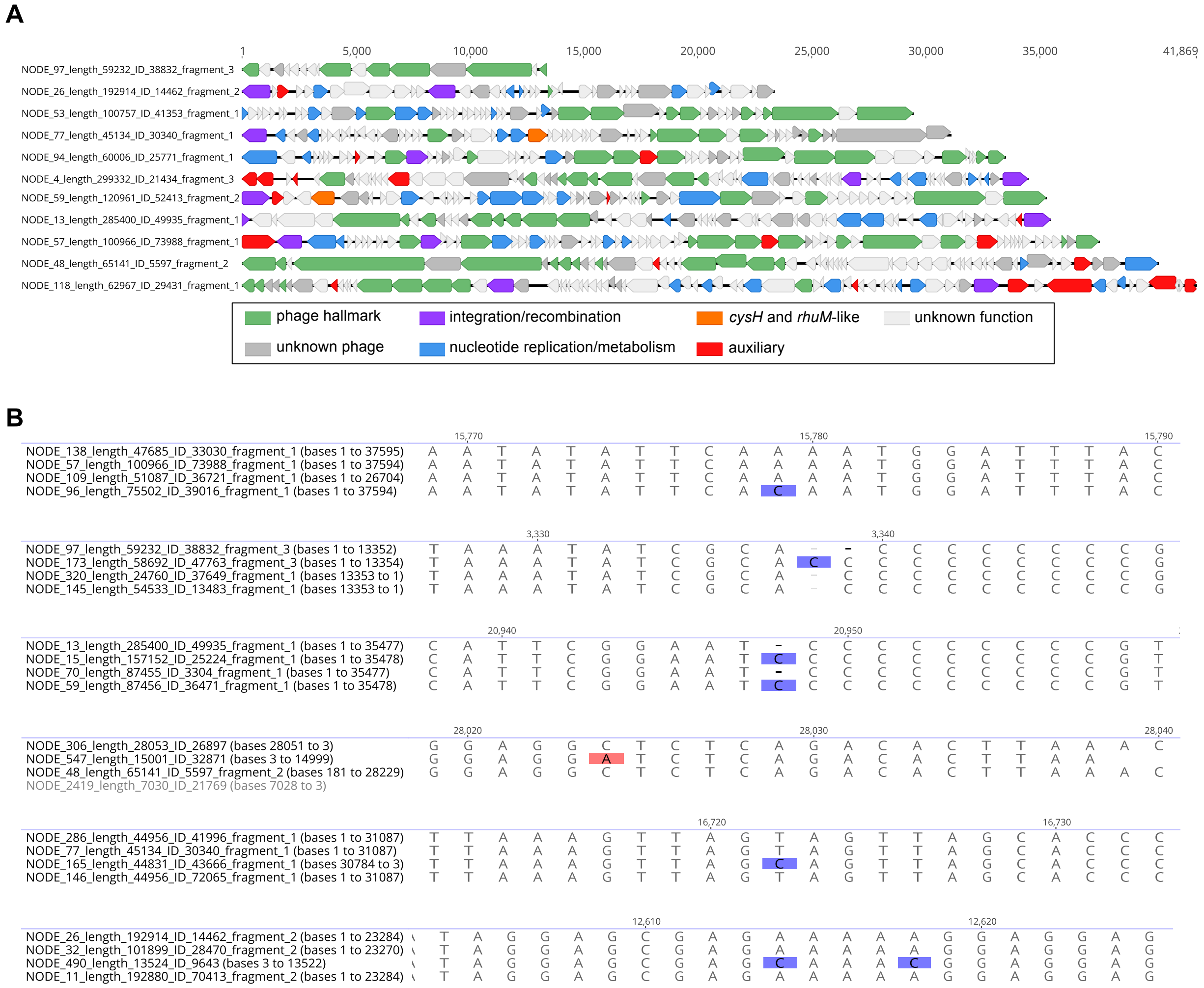


**Supplemental Figure S2**. **Eleven prophage representatives dormant over time**. (A) Genome diagrams and annotations of 11 human gut prophages identified to be dormant over multiple time points. The longest representative from each population was selected for visualization. (B) Regions of dissimilarity in genome alignments per population.

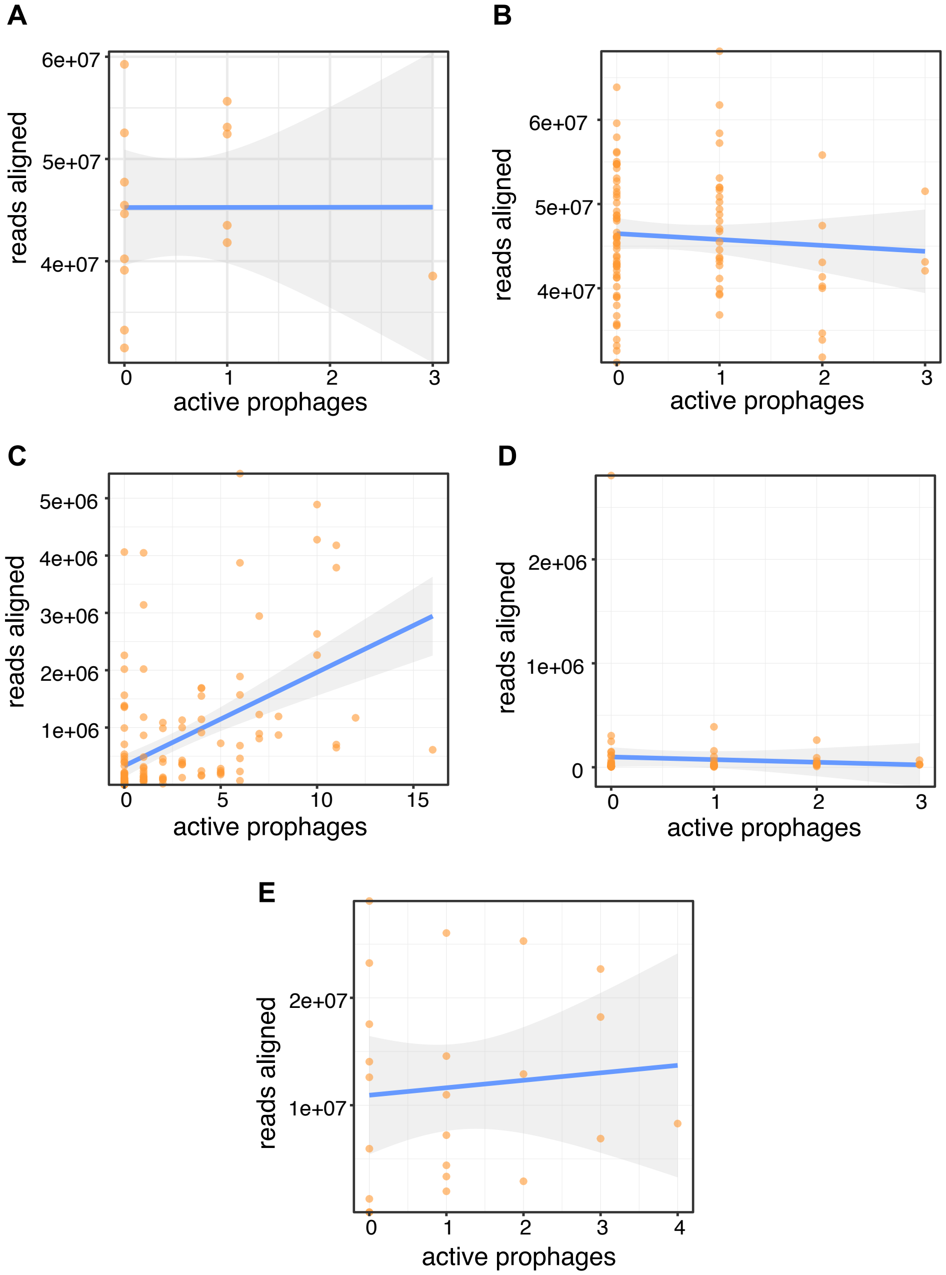


**Supplemental Figure S3. Correlation of total reads aligned to total active prophages identified.** Each plot represents a metagenomic set corresponding to **Table 1** (A:a, B:b, C:c, D:d and E:e). Each dot represents a single metagenomic sample. Linear regression lines are shown in blue with 95% confidence intervals shaded in gray.

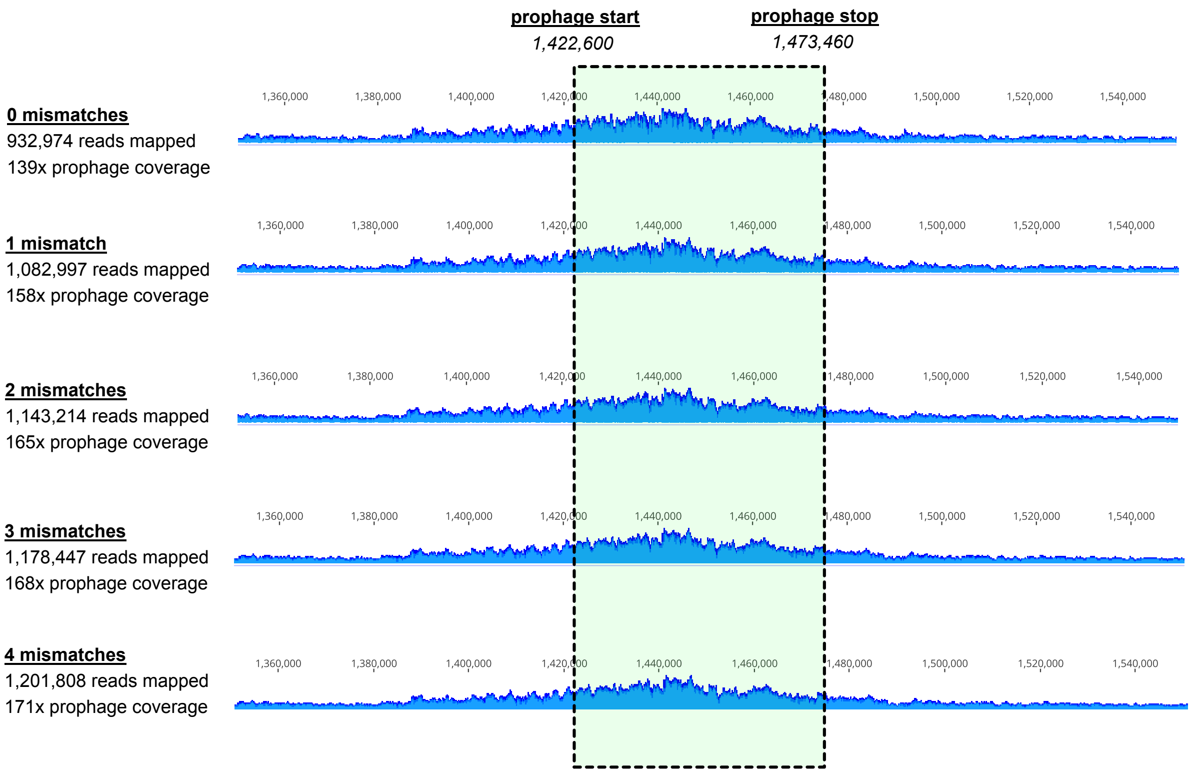


**Supplemental Figure S4. Benchmarking of read alignment maximum mismatches threshold**. An example active *Bacillus licheniformis* DSM13 prophage was assessed by PropagAtE with the maximum allowed mismatches set from 0-4. Each yielded relative results except for total reads mapped and average prophage coverage.
